# Foliar pathogen and drought impose reproducible but community-dependent influence on the root microbiome

**DOI:** 10.64898/2026.09.02.748968

**Authors:** Edda Francomano, Meriem Miyassa Aci, Nesma Zakaria Mohamed, Leonardo Schena, Antonino Malacrinò

## Abstract

Plant microbiomes are assembled from environmental pools that differ substantially in composition, yet whether their responses to stress follow general rules or depend on the resident community remains unclear. We tested the generality of root microbiome responses to stress by growing three tomato (*Solanum lycopersicum*) genotypes in 20 independently sourced microbial communities under controlled abiotic and biotic stress. Microbial communities were transferred into a common sterile substrate, allowing to vary microbial community identity independently of soil physicochemical properties, and plants were exposed to drought, the foliar pathogen *Pseudomonas syringae* pv. *tomato*, or no stress. We analyzed the root microbiota using 16S rRNA amplicon sequencing. Stressor identity explained more variation in bacterial community composition than inoculum identity or plant genotype. Hierarchical models showed that many bacterial taxa responded consistently across distinct starting communities, with taxon identity contributing far more variation in stress response than microbial community of origin. Both stressors shifted between-community dissimilarity from taxon turnover toward nestedness, indicating increasingly similar patterns of taxon loss, but the pathogen produced stronger effects and increased variability among replicate plants. Drought reduced phylogenetic redundancy, whereas pathogen increased dispersal limitation. Together, these results show that stress imposes reproducible ecological filters across diverse starting microbiomes, while the magnitude and resulting community state remain contingent on the resident microbial community. This provides a basis for identifying transferable microbial targets for microbiome-based crop stress management, while also managing the local microbiome to maximize their beneficial effects.

## Introduction

Plant roots are colonized by microbial communities recruited largely from the surrounding soil (Mohamed *et al*. 2024; Trivedi *et al*. 2020), and the composition of these communities affects plant fitness, nutrition, and health (Francomano *et al*. 2026b; Trivedi *et al*. 2020). Soils differ in the microbial taxa they contain, and individual plants of the same genotype growing on soil from different locations recruit root communities with partial taxonomic overlap (Beschoren da Costa *et al*. 2022; Edwards *et al*. 2023; Simonin *et al*. 2020). At the same time, when the plant holobiont experiences stress, it reshapes its microbiome, recruiting microorganisms from the surrounding soil to help counteracting the stress (Francomano *et al*. 2026b; Rolfe *et al*. 2019; Trivedi *et al*. 2020). This has been shown to occur in response to herbivory (Malacrinò *et al*. 2021b; Malacrinò & Bennett 2024; Rodríguez-Blanco *et al*. 2026), pathogens (Alfaro-García *et al*. 2025; Francomano *et al*. 2026a; Mendes *et al*. 2023), drought or heat (Laine & Leino 2025; Pantigoso *et al*. 2025; Tiziani *et al*. 2022), and several other stressors. Yet, we still know little on whether the root microbiota is restructured following generalized rules or whether this response depends on the resident soil community. This information is key to any attempt to microbiome-management practices.

Soil is consistently identified as the dominant source of variation in root community composition, with host genotype and other factors accounting for smaller fractions (He *et al*. 2024; Malacrinò *et al*. 2021a; Malacrinò & Bennett 2024). Both biotic and abiotic stressors influence soil- and plant-associated microbial communities (Hopkins *et al*. 2025; Li *et al*. 2022; Liu *et al*. 2023; Wang *et al*. 2025; Xiong *et al*. 2026). Different stressors act through different mechanisms. Drought, for example, acts on both soil and plant at the same time, with direct and indirect (plant-mediated) effects on both plant and soil microbiomes (Schimel 2018; Trivedi *et al*. 2022). Drought has repeatedly been reported to enrich Actinomycetota in the roots and rhizosphere (Cosma & Abeel 2026; Fonseca-Garcia *et al*. 2025; Swift *et al*. 2024). Pathogen infection also alters root and rhizosphere communities, even when infection is confined to the leaves (Becker *et al*. 2023; Luo *et al*. 2022, 2025). In foliar infections, for example, the pathogen is confined to leaf tissue, the substrate is unaltered, and the only available route is the plant immune signaling, altering root exudation and the recruited microbial community (Yuan *et al*. 2018). Almost all of these studies, however, are performed on a single soil, or a small number of soils, while studies where multiple soils have been compared (Guo *et al*. 2024; Taketani *et al*. 2026; Walsh *et al*. 2021) do not explore the influence of stressors on those microbial communities. Thus, we still lack a generalized understanding of how stressors influence the root microbiome, which ecological processes are involved, and which patterns generalize across soil communities.

In this study, we aim to fill this knowledge gap and generalize knowledge about the root microbiome assembly under stress by growing three different genotypes of tomato seedlings (*Solanum lycopersicum* L.) on soils hosting 20 different microbial communities, and exposing them to two stressors: drought and the pathogen *Pseudomonas syringae* pv. *tomato*. We hypothesized that stress responses of the root microbiome generalize across soil communities, while maintaining a stress-specific signature, and we tested four main predictions. First, stressor identity should account for more variation in root microbiota composition than the identity of the soil community. Second, a set of taxa should respond to a stressor in the same direction across soil communities. Third, if stress exerts selective pressure, microbiota dissimilarity should shift from turnover towards nestedness, and replicate plants should become less similar to one another as the outcome of selection becomes more variable. Fourth, if stress alters the balance of assembly processes, as it has been reported along stress gradients in soil (Ning *et al*. 2020), the change should appear as a shift between deterministic and stochastic processes that is consistent across soil communities. Together, testing these predictions provides the first empirical evidence of generalized ecological effects of major stressors on the plant microbiome, which is key in designing microbiome-based strategies to reduce their impact on our crops.

## Results

In this study we implemented a design in which the source of the microbial community was varied while other factors were held constant. Microbial communities were extracted by centrifugation from 20 soils and inoculated into a common autoclaved substrate, so that the soils contributed their microbiota but not their physical or chemical properties. The applied communities differed substantially from one another, as source identity explained 69.2% of compositional variation among the inoculum libraries (PERMANOVA on Aitchison distance, F = 3.03, p = 0.001), and replicate extractions of the same soil were markedly less dissimilar than extractions of different soils (p = 0.001 by permutation), and a median of 26% of the ASVs detected in a source were found in no other inoculum. Three tomato genotypes were grown in each community and exposed to drought, imposed by withholding water until plants wilted, to the foliar pathogen *Pseudomonas syringae* pv. *tomato* DC3000, or to no stressor. Root bacterial communities were characterized using 16S rRNA gene amplicon sequencing and analyzed in a compositional framework, with inference from Bayesian multilevel models.

### Stressor drives the structure of the root microbiome

Permutational analysis of Aitchison distances identified significant but small effects driven by each factor when fitted marginally: soil inoculum R^2^ = 0.041 (F_19, 942_ = 2.12, p = 0.001), stressor R^2^= 0.027 (F_2, 959_ = 13.35, p = 0.001) and genotype R^2^= 0.010 (F_2, 959_ = 4.76, p = 0.001; **Fig. 1A,B, Supplementary Table S1**). In the full model the largest single term was the inoculum × stressor interaction (R^2^= 0.073, F_38, 858_ = 2.08, p = 0.001; **Supplementary Table S1**). PERMANOVA R^2^ cannot be compared across terms of very different degrees of freedom, so we decomposed variance in a Bayesian multilevel model of the five leading Aitchison principal components. Stressor accounted for the largest share (0.297, 95% credible interval [CrI] 0.129-0.506), followed by the inoculum × stressor interaction (0.130 [0.086-0.178]) and genotype (0.088 [0.016-0.241]; **Fig. 1C, Supplementary Table S2**). Together, this first analysis suggests that soil source strongly conditioned how that community responded to stress.

**Figure 1.**
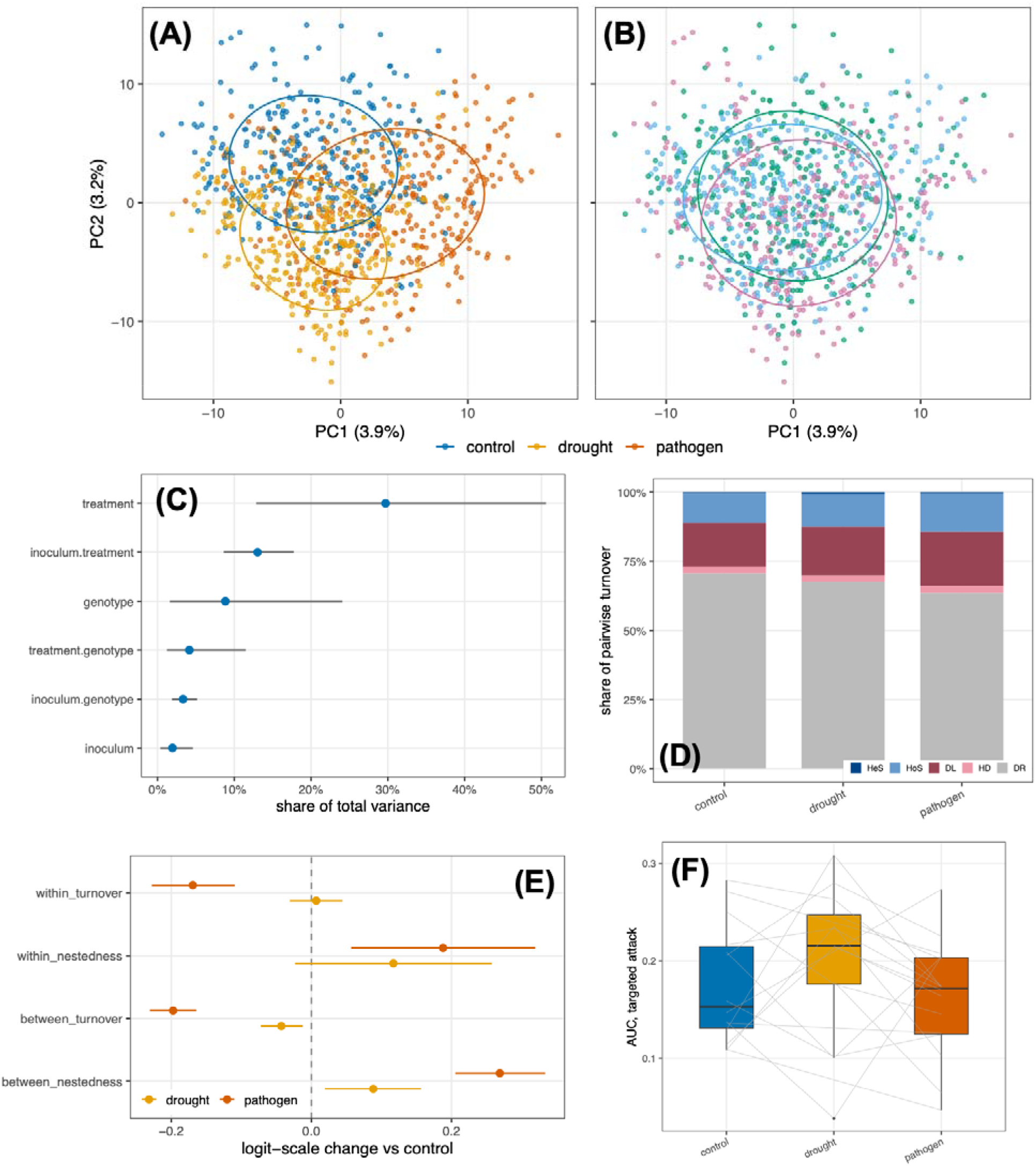
**(A)** Principal component analysis of centered log-ratio-transformed ASV abundances (Aitchison ordination) colored by stressor, with ellipses enclosing 68% of samples per group. **(B)** The same ordination colored by tomato genotype. **(C)** Share of variance attributable to each design factor, from a Bayesian multilevel model of PC1-PC5 weighted by the Aitchison variance each component carries; points in are posterior means and bars 95% credible intervals. Note that PC1-PC5 together carry 12.0% of total Aitchison variance. **(D)** Mean share of pairwise turnover assigned by iCAMP to each of five processes, by stressor. **(E)** Posterior stressor effects on each component of Sørensen dissimilarity partitioned into turnover and nestedness from Bayesian beta regression, expressed on the logit scale relative to control. **(F)** Robustness of co-occurrence networks under targeted attack by stressor; grey lines join networks built from the same soil source.

It is important to note that multivariate dispersion was heterogeneous among stressors (F_2, 959_ = 83.35, p = 0.001), inocula (F_19, 942_ = 3.31, p = 0.001), genotypes (F_2, 959_ = 3.02, p = 0.041). Part of the stressor signal in the PERMANOVA is therefore a difference in data dispersion but this result is caused by an increased variability in microbiome composition after stress. Indeed, replicated plants within the same treatment differed more from one another under the pathogen (+0.137 [0.086, 0.186]) but not under drought (+0.034 [−0.004, 0.071]; **Supplementary Table S3**). Distance between different soil communities increased under both stressors, though more strongly under the pathogen (+0.148 [0.131, 0.165]) than under drought (+0.029 [0.014, 0.044]; **Supplementary Table S3**). Slopes for individual inocula showed the pathogen effect was general, being credible in 16 of 20 inocula, against 9 of 20 for drought (**Supplementary Fig. S1**).

Partitioning Sørensen dissimilarity showed that between soil communities, community turnover decreased under both pathogen (−0.197 [−0.231, −0.164]) and under drought (−0.043 [−0.072, −0.012]), while nestedness increased (pathogen +0.269 [0.205, 0.334]; drought +0.088 [0.019, 0.157]; **Fig. 1E, Supplementary Table S4**). The pathogen effect on both components was credible in all 20 inocula, the drought effect in 6 of 20 for turnover and 2 of 20 for nestedness (**Supplementary Fig. S2**). Phylogenetic redundancy declined under drought (−0.035 [−0.056, −0.013]) but showed no credible response to the pathogen (+0.016 [−0.013, 0.046]; **Supplementary Table S5**). Per-inoculum slopes were credible in 7 of 20 inocula for drought and 6 of 20 for the pathogen, indicating a modest and context-dependent effect (**Supplementary Fig. S3**).

Together, these results indicate that the two stressors restructured the root microbiome in different ways. The pathogen made replicate communities less similar to one another and shifted the difference between soil communities from turnover towards nestedness, an effect credible in nearly every inoculum, while drought produced the same qualitative shift but more weakly, and instead reduced the phylogenetic redundancy.

### Assembly is dominated by drift, and the pathogen shifts it towards dispersal limitation

Across 18,676 sample pairs within 20 soil communities, drift and other stochastic processes accounted for 64-71% of turnover, dispersal limitation for 16-20%, and selection for only 11-14% (homogeneous 10.6-13.7%; heterogeneous 0.5-0.6%; **Fig. 1D**). The pathogen credibly increased the proportion of deterministic processes (+0.300 [0.071, 0.536]), whereas drought did not (+0.118 [−0.102, 0.339]), and per-source slopes were credible in 10 of 20 inocula for the pathogen and 8 of 20 for drought (**Supplementary Fig. S4-S5, Supplementary Table S6**). Within the stochastic component, exposure to the pathogen increased dispersal limitation (+0.266 [0.089, 0.442]) and decreased drift (−0.339 [−0.456, −0.217]), and drought also decreased drift (−0.148 [−0.287, −0.010]) without a credible effect on dispersal limitation (+0.148 [−0.070, 0.369]) (**Supplementary Fig. S4-S5, Supplementary Table S6**).

We inferred also microbial co-occurrence networks and found that robustness to targeted node removal varied among soil sources (area under the attack curve 0.064-0.257, mean 0.129±0.050) and was lower under random removal (0.306±0.053; **Supplementary Table S7**). This variation was, however, almost entirely explained by network size, as AUC correlated with edge count at r = 0.952 and with modularity at r = −0.873. Once edge count was included as a covariate, we did not observe any effect of stressor on network robustness (drought −0.090 [−0.207, 0.029]; pathogen −0.106 [−0.215, 0.008]; **Fig. 1F, Supplementary Table S8**). We therefore find no evidence that either stressor changed network robustness.

Together, these results indicate that stress shifted assembly modestly towards deterministic processes under the pathogen alone, and redistributed the stochastic component itself, replacing drift with dispersal limitation. The robustness of the co-occurrence network of the community was unaffected by either stressor.

### A reproducible set of taxa responds to each stressor across soil communities

We identified patterns in ASV variation under stress by pooling across soil sources in a hierarchical meta-analysis, and found that out of the 150 most consistently responding ASVs per stressor, 108 had a credibly non-zero pooled effect under drought and 106 under the pathogen, of which 94 and 86 respectively also agreed in direction in at least 70% of the inocula in which they were testable (**Supplementary Table S9**). Under drought (*n* = 45 enriched, *n* = 49 depleted), Proteobacteria dominated (*n* = 47 of 94) followed by Actinobacteriota (*n* = 15; **Fig. 2, Supplementary Table S9**). The main enriched ASVs included *Microbacterium, Pseudomonas, Brevundimonas* and *Arthrobacter*, while those reduced under drought stress included *Haliangium*, rhizobia, Gemmatimonas and *Sandaracinus*. In plants exposed to the pathogen (*n* = 46 enriched, *n* = 40 depleted; **Fig. 2, Supplementary Table S9**), enriched ASVs included *Rhodobacter, Jahnella, Arthrobacter* and *Micrococcus*, while ASVs identified as rhizobia, Caulobacteraceae and *Shinella* decreased in their relative abundance. Thirty-eight taxa were general responders to both stressors and 37 of these moved in the same direction (**Supplementary Table S9**), with pooled effects correlated at r = 0.726. Several ASVs identified as rhizobia were consistently depleted under drought, and were the most strongly depleted taxa under the pathogen as well, although one of the four rhizobia ASVs responding to the pathogen was instead enriched.

**Figure 2.**
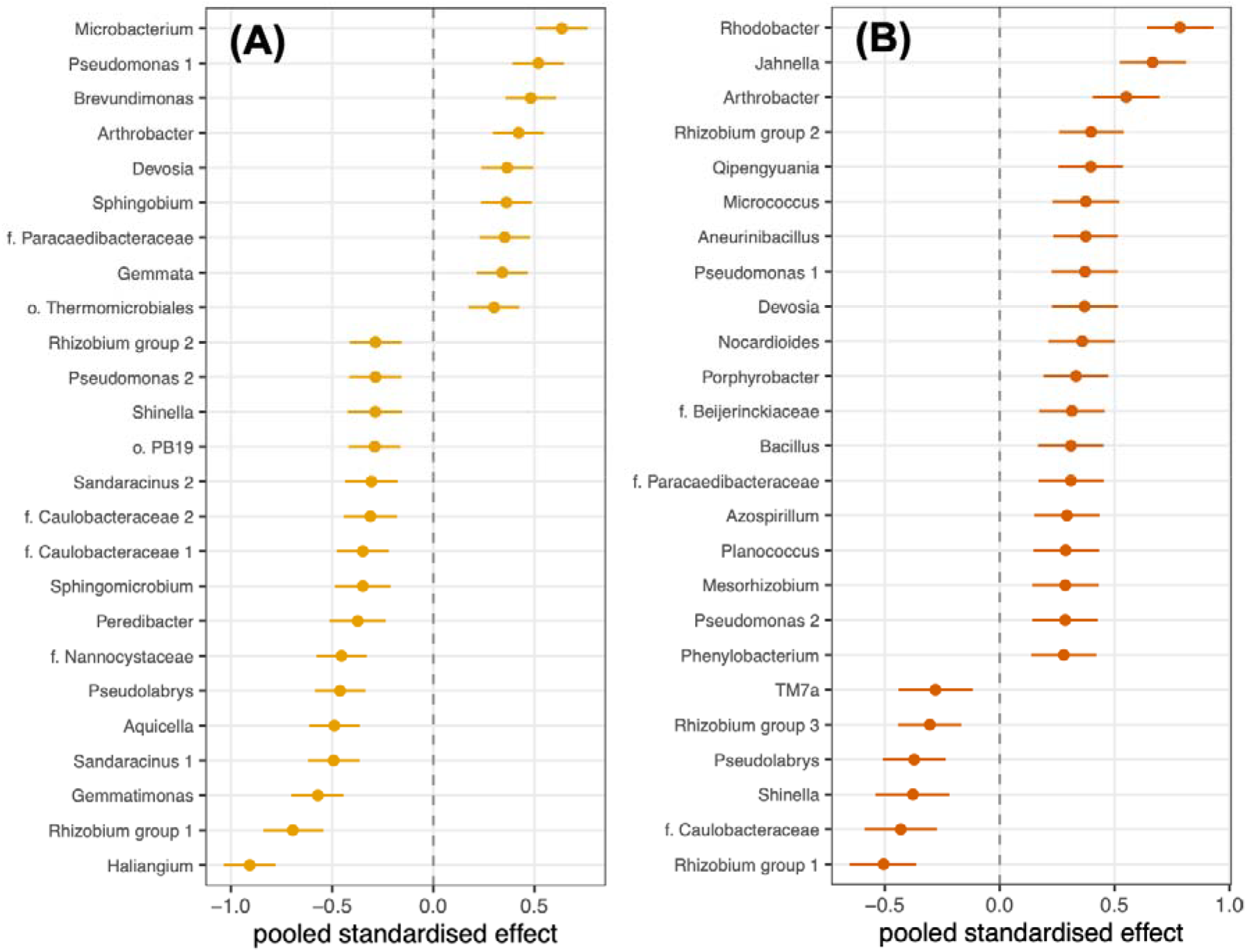
A reproducible set of taxa responds to each stressor across soil communities. **(A, B)** Pooled posterior effects of **(A)** drought and **(B)** the pathogen for the taxa meeting both the credibility and the directional-consistency criteria, from a hierarchical meta-analysis across soil sources; bars are 95% credible intervals.

**Figure 3.**
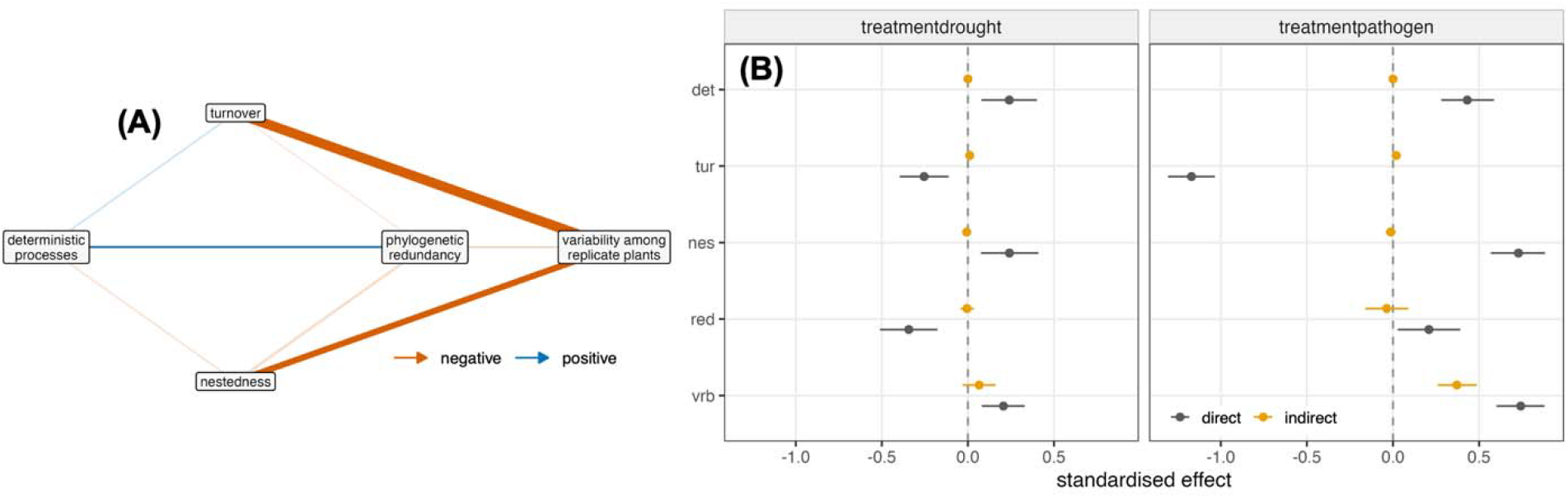
A path model locates the mechanism in dissimilarity structure. **(A)** Structural equation model of the mediation chain; arrow width is the absolute standardized path coefficient, color its sign, and faded arrows are paths whose 95% credible interval includes zero. **(B)** Direct and summed indirect components of each stressor effect. All variables were put on an unbounded scale and standardized so that path coefficients are comparable and their products are interpretable as indirect effects; magnitudes are therefore not comparable with the single-equation estimates reported elsewhere. det = deterministic processes; tur = turnover; nes = nestedness; red = phylogenetic redundancy; vrb = variability among replicated plants.

This model separates three sources of variation: differences between taxa in how strongly they respond, differences between inocula in the average response they support, and the extent to which a given taxon responds differently depending on the inoculum it is measured in. Because each ASV is estimated once per inoculum, this last term is the residual. Between-taxon variation was 0.252 [0.223, 0.285] under drought and 0.247 [0.218, 0.278] under the pathogen, taxon×inoculum variation 0.298 [0.290, 0.306] and 0.337 [0.328, 0.346], and between-inoculum variation 0.011 [0.001, 0.028] and 0.025 [0.002, 0.062] (**Supplementary Table S10**). The taxon×inoculum variance was comparable to the between-taxon variance, whereas the between-inoculum variance was an order of magnitude smaller. Stress responses were therefore taxon-specific and inoculum-dependent, but no inoculum conferred a systematically stronger or weaker response.

Together, these results indicate that whether a taxon responds to stress is determined more by its identity than by the community of origin. The substantial overlap between the two stressor responses, and the opposing responses of rhizobia and Actinobacteria in both, point to a shared component of the root community’s response to stress.

### A path model identifies the mechanism in dissimilarity structure, not in assembly process

Finally, we combined all the metrics above in a Bayesian structural equation model (856 samples, 89.0% of the compositional dataset, 20 inocula). The model explained 48.4% of variance in variability among replicate plants, 34.8% in turnover, 11.5% in nestedness, 12.3% in redundancy and 11.9% in assembly deterministic processes (**Supplementary Table S11**). Both stressors did increase the proportion of deterministic processes (drought +0.240 [0.078, 0.400]; pathogen +0.431 [0.281, 0.587]), but deterministic processes had no credible onward effect on turnover (+0.043 [−0.014, 0.101]), nestedness (−0.030 [−0.097, 0.037]) or variability (−0.005 [−0.058, 0.049]), with only a weak positive downstream path to phylogenetic redundancy (+0.070 [0.001, 0.139]; **Fig. 4, Supplementary Table S12**). Instead, the stressors acted on dissimilarity directly. The pathogen strongly reduced turnover (−1.171 [−1.307, −1.035]) and increased nestedness (+0.729 [0.568, 0.883]), and both turnover (−0.520 [−0.595, −0.447]) and nestedness (−0.309 [−0.371, −0.247]) predicted lower variability, whereas redundancy did not (−0.032 [−0.084, 0.021]) (**Fig. 4, Supplementary Table S12**). Decomposing the pathogen total effect on variability (1.112 [0.972, 1.255]) estimated a credible indirect component of 0.371 [0.260, 0.487] mediated through the chain. For drought the summed indirect effect was not credible (0.065 [−0.030, 0.161]) although individual routes were, in opposing directions: via turnover +0.133 [0.057, 0.212] and via nestedness −0.075 [−0.131, −0.023] (**Fig. 4, Supplementary Table S12**).

Together, these results indicate that the stressors did not act on the root community through the assembly processes they altered, since the shift towards deterministic processes had no consequence downstream. The effect ran instead through the structure of dissimilarity, and about a third of the pathogen effect on variability was transmitted this way, the remainder acting directly.

## Discussion

In this study, we grew three tomato genotypes in twenty independently sourced microbial communities and exposed them to drought or to a foliar pathogen and tested whether stressors produce generalized effects across soil communities and plant genotypes. Stressor identity accounted for more variation in root community composition than other factors. A subset of taxa responded to a stressor in the same direction across communities. Both stressors shifted the difference between communities from turnover (replacement of taxa) to nestedness (loss of taxa), and made replicate plants less similar to one another. Finally, a path model showed that the stressors acted through the structure of microbiota dissimilarity rather than through the assembly processes they also altered.

Stressor identity explained the largest portion of variation among the main compositional axes, followed by the interaction between stressor and source community, and the source community identity. Previous studies reported soil effects spanning a wide range of explained microbiota variation, from 70.6% of rhizosphere variation in soybean grown in agricultural versus forest soil (Liu *et al*. 2019) and 19.7% of root endosphere variation in tomato grown with three distinct soil communities (Malacrinò & Bennett 2024), to 8.6% in chrysanthemum roots (Pangesti *et al*. 2020) and roughly 3% in *Medicago truncatula* (Brown *et al*. 2020). While in this study we observed a smaller amount of variation explained by inoculum source compared to our previous results on a similar system (Malacrinò & Bennett 2024), here we used a different inoculation technique, which might have influenced the results (Howard *et al*. 2017) although this choice was motivated by experimental reasons to control for soil abiotic conditions and accurately test our hypotheses. Similar range of variation was reported for the portion of microbiota variance explained by a stressor. In a previous study, foliar pathogen infection explained 19.5% of variation in apple root communities in a two-level contrast with ten plants per group (Becker *et al*. 2023), whereas drought explained 5.9% of rice root endosphere variation across three soils and four genotypes (Santos-Medellín *et al*. 2017) and 3.0% in *Arabidopsis* (Wang *et al*. 2024). It is important to note, however, that PERMANOVA R^2^ is not comparable across studies of different size, factor structure and heterogeneity, and these comparisons should be treated as explorative.

Replicate plants sharing a source community and genotype differed more from one another under stress, and this effect was larger for the pathogen treatment than drought. Increased among-individual variation under stress supports the argument that perturbation reduces the capacity of the host or its microbiota to regulate community composition, so that stressed individuals diverge from one another rather than converging on a new state (Arnault *et al*. 2022). Because the pathogen was confined to the leaves and the substrate was untreated, the increase in among-plant variation cannot be a direct effect on the microorganisms and it likely has been transmitted through the host. The most likely route is that foliar infections alter root exudation and thereby the community recruited to the root (Berendsen *et al*. 2018; Yuan *et al*. 2018). For the host, this could mean that the root community recruitment is less predictable under stress, and any benefit that depends on a particular community composition becomes less reliably available. On the other hand, this study focuses on the community taxonomical composition, and it does not infer function. Thus, while the recruited community becomes more taxonomically variable under stress, it might still recruit taxa with similar functional role. Our results suggest that the stressors acted as a common filter, and the more strongly a community was filtered the more it came to resemble a depleted subset of the others. This is consistent with the selective enrichment of stress-tolerant lineages reported under drought across many host species (Cosma & Abeel 2026; Naylor *et al*. 2017; Xu *et al*. 2018). Interestingly, this selective enrichment might remove taxa non-randomly and leave different survivors in different individuals, increasing variation across them.

Phylogenetic redundancy declined under drought but not under the pathogen treatment. A decline in redundancy, under the insurance hypothesis (Barnett & Shade 2025; Ramond *et al*. 2025; Yachi & Loreau 1999), may indicate that drought removed taxa in a way that concentrated loss within lineages rather than spreading it across them, potentially leaving the community with a restricted functional buffer. That the pathogen did not produce the same decline, despite producing much larger effects on other metrics, suggests drought might have selected microbes on traits that are phylogenetically conserved (Metze *et al*. 2023), and the pathogen on traits distributed more evenly across the phylogeny. Thus, a root microbiota influenced by drought may be less able to withstand a subsequent perturbation than one influenced by a pathogen. Again, as above, these are speculative assumptions as this study inferred the community taxonomical composition and not function, and further tests are needed to clarify this result.

Drift and other stochastic processes accounted for roughly two thirds of turnover between plants, followed by dispersal limitation and selection. Dominance of stochastic processes in root-associated communities has been reported repeatedly (Ai *et al*. 2026; Francomano *et al*. 2026a; Li & Gao 2023; Li *et al*. 2025; Mosca *et al*. 2026). Neither stressor changed the balance between deterministic and stochastic processes. The pathogen exposure increased the proportion of dispersal limitation and decreased drift. We expected any shift in dispersal limitation to accompany drought, because only drought alters the physical medium in which dispersal occurs, as reduction in soil water content thins and disconnects the water films that occupy soil pores and so restricts bacterial movement (Malik & Bouskill 2022; Schimel 2018). A foliar pathogen leaves the substrate unaltered, so an increase in dispersal limitation cannot be a physical effect and must reflect the root environment itself, a new direction worth of additional testing, since it implies that host-mediated changes can constrain microbial dispersal without any change in soil.

Robustness to targeted node removal varied among source communities, but almost all of that variation was explained by the number of edges recovered, and once edge count was accounted for neither stressor altered it. Comparisons of network robustness between treatments are common in the microbiome literature and are frequently made without this control, even though network size and density are known to determine most other network statistics (Connor *et al*. 2017). We found no evidence that either stressor changed the co-occurrence structure of the root community beyond what its effect on the number of detectable associations implies. Thus, either the co-occurrence structure did not change, or networks inferred from tens of samples over hundreds of taxa are too sparse to detect a change of the size present.

Pooling across source communities in a hierarchical model identified taxa whose response to a stressor was both credible and directionally consistent across the communities in which they could be tested, and the variance attributable to taxon identity was several times that attributable to the community of origin. Drought enriched Actinomycetota including *Microbacterium, Arthrobacter* and *Nocardioides*, matching the enrichment reported in the roots of grasses (Naylor *et al*. 2017), sorghum (Xu *et al*. 2018) and rice (Santos-Medellín *et al*. 2021), and generalized across hosts in a recent meta-analysis (Cosma & Abeel 2026). Foliar pathogen enriched a partly overlapping set, consistent with the recruitment of specific bacterial taxa to the rhizosphere following foliar defense activation (Berendsen *et al*. 2018) and with the exudate-mediated legacy of aboveground infection (Yuan *et al*. 2018). The consistent depletion of taxa identified as rhizobia under both stressors, as far as we are aware, has not been reported as a general stress response. We also found that the response of a taxon to stress is largely a property of that taxon, and therefore that a taxon identified as stress-responsive in one soil may be a reasonable candidate in another. This is a key assumption for microbiome-based management of crop stress, and it has not previously been tested across this many independent communities, although this needs to be further evaluated under field conditions and across different soil types.

The analyses above treated each community property separately. Using a path model, we asked whether the stressors changed one metric because they changed another. We had expected a chain in which stress strengthens selection, stronger selection changes which taxa differ between communities, and that in turn changes how reproducible the assembled community is. Both stressors increased the fraction of turnover attributable to deterministic processes, the pathogen more than drought. However, that fraction had no credible effect on turnover, on nestedness, or on variability among replicate plants. Selection strength responded to stress and did not influence other metrics. What carried the effect was the composition of the dissimilarity itself. The pathogen sharply reduced turnover and increased nestedness, thus communities came to differ less in which taxa they held and more in how many they had lost, and both predicted lower variability among replicate plants.

In this study, stressor identity outweighed source community identity in explaining composition, the same taxa responded in the same direction across communities, and both stressors shifted communities from differing by replacement to differing by loss, with the pathogen making the microbiota of replicate plants less similar to one another, and drought reducing its phylogenetic redundancy. The finding that a foliar pathogen restructures a root community it never contacts, alters the dispersal component of that community assembly, and does so consistently across different starting communities, suggests that the host is imposing a filter whose consequences are reproducible across communities. The generality of the taxon-level responses is key to manage crop stress through microbiome-based solutions, and it depends on the assumption that a taxon identified as beneficial or stress-responsive in one location will act comparably in another, and this assumption has rarely been tested against more than a handful of soils. Our data support it for the direction of response, while making clear that its magnitude depends on the whole microbial community. Together, these results suggest that stressors can impose reproducible taxonomic changes across diverse starting communities, but the extent of community restructuring and the resulting ecological state remain contingent on the initial microbiome. This distinction will be important for developing microbiome-based approaches to crop stress, because predictable taxon-level responses may provide transferable targets, and the management of the local microbiota can modulate their ability of being recruited and exert beneficial effects.

## Methods

### Experimental design

The experiment tested how the root bacterial community assembles from a defined microbial source, and included 20 microbial inocula, 3 levels of stressors (drought, pathogen, control), 3 plant genotypes, in a full factorial design replicated over 7 plants per group, for a total of 1,350 plants.

Seeds of three tomato (*Solanum lycopersicum* L.) genotypes (G1: Moneymaker, G2: Cuor di Bue beefsteak tomato, and G3: Trixi KS cherry tomato; Sativa Biosaatgut GmbH, Germany) were germinated on small pots (5 cm height, 3 cm basal diameter). All plants were grown in a common background substrate of sand and sieved soil (2:1 v/v; autoclaved twice for 3h at 121°C), so that experimental units differed in the microbial communities introduced and not in the physical or chemical properties of the growing medium. Plants were grown under full-spectrum artificial lighting on a 16:8 h light:dark cycle.

Twenty microbial inocula were prepared from soils collected at sites differing in land use, and soil type, including cultivated land under several management regimes, forest and uncultivated land (**Table S13**), chosen to maximize variation in the communities introduced. Inocula were extracted by a washing procedure (Mohamed *et al*. 2024; Walsh *et al*. 2021): soil was vortexed in sterile distilled water, centrifuged to remove coarse particles, and the microbial pellet resuspended in sterile phosphate-buffered saline (PBS). Each plant received 1 mL of a single inoculum.

Plants were assigned to one of three treatments: an unstressed control, drought, or exposure to the pathogen *Pseudomonas syringae* pv. *tomato* (*Psto*) DC3000. Plants were exposed to stressors at 6 weeks after germination for 30 days. Drought was imposed by withholding water until plants approached wilting. Control plants were watered normally throughout. *Psto* was applied once, as an aerosol delivered to the foliage, at 10^8^ CFU/mL. Because the pathogen was applied to the leaves and the substrate was not treated, the pathogen did not contact the root community directly. At the end of the treatment period, root samples were collected from each plant, frozen immediately and stored at −80°C, then freeze-dried before nucleic acid extraction. Samples were not surface-sterilized, so the communities characterized include both epiphytic and endophytic taxa.

### DNA extraction, library prep, and amplicon sequencing

DNA from individual samples was extracted using the DNeasy PowerMax Soil Kit (Qiagen), and successful extraction was verified using a Nanodrop 2000 (Thermo Scientific, USA) spectrophotometer. Bacterial communities were characterized by amplifying the 16S rRNA gene with the primer pair 515F/806R (Caporaso *et al*. 2012). PCR was carried out in 25 µL containing approximately 50 ng template DNA, 0.5 µM of each primer, 1X KAPA HiFi HotStart ReadyMix (Roche, USA) and nuclease-free water, in a Mastercycler Ep Gradient S (Eppendorf, Germany) under the following conditions: 95°C for 3 min; 35 cycles of 98°C for 30 s, 55°C for 30 s and 72°C for 30 s; and a final extension at 72°C for 10 min. Non-template controls, in which the sample was replaced with nuclease-free water, were amplified alongside every plate to account for contamination of instruments, reagents and consumables. Products were purified (Agencourt AMPure XP, Beckman Coulter), used in a second short PCR to attach Illumina adaptors, and purified again. Libraries were quantified using a Qubit fluorometer (Thermo Fisher Scientific, USA), pooled at equimolar concentrations, and their fragment size was verified using a TapeStation 4150 system (Agilent Technologies, USA). Sequencing was performed on an Illumina NovaSeq platform (Illumina, USA) using a SP flow cell and 2 × 250bp paired-end chemistry.

### Data processing and analysis

Raw sequencing reads were processed using the *nf-core/ampliseq* pipeline v2.7.1 (Ewels *et al*. 2020; Straub *et al*. 2020). Briefly, reads were quality filtered, primer sequences were removed, amplicon sequence variants (ASVs) were inferred using DADA2 (Callahan *et al*. 2016), and chimeric sequences were identified and discarded. Taxonomic assignment was performed against the SILVA database v138 (Quast *et al*. 2013). Representative bacterial ASV sequences were aligned using *MAFFT* v7.525 (Katoh *et al*. 2002), and a phylogenetic tree was inferred with *FastTree* v2.1.11 (Price *et al*. 2009). All downstream processing was conducted in *R* v4.4.1 (R Core Team 2020). Sequencing yielded 1,309 bacterial libraries containing 12,962 ASVs and 19.6 million reads.

ASV tables, taxonomic assignments, sample metadata, and phylogenetic trees were imported into *phyloseq* v1.48 (McMurdie & Holmes 2013). Non-target sequences were removed first: ASVs not assigned to Bacteria or Archaea, and those assigned to the order Chloroplast or the family Mitochondria, which derive from the host. Although these accounted for only 122 ASVs, they carried 19.5% of all reads. Libraries with fewer than 3,000 reads after this step were discarded. Non-template controls were exempt from this threshold, as they were used to identify potential contaminant ASVs using the prevalence-based method implemented in *decontam* v1.24 (Davis *et al*. 2018). Because each PCR plate carried its own non-template controls, the model was fitted separately for each plate and an ASV was called a contaminant if it was called on any plate. Five ASVs were identified and removed, accounting for 4.5% of reads. ASVs occurring in fewer than 5% of root samples, or with fewer than five reads in total, were removed, retaining 494 ASVs and 90.6% of reads. Analyses requiring presence-absence or abundance-weighted data, rather than log-ratios, were performed on a table rarefied to the 10th percentile of library size.

Of the libraries sequenced, 1,024 root samples passed quality control. The loss was distributed across the design as we retained data from 178 of the 180 combinations of inoculum, stressor and genotype, 170 of these retained at least three plants, and the median combination retained six (**Supplementary Fig. S6**). Retaining three genotypes across all 20 soil communities meant that each stressor contrast was replicated many times over, and because every effect was estimated separately within each soil community and then pooled in a Bayesian multilevel model, combinations with fewer plants contribute less to the pooled estimate rather than being weighted equally or discarded.

ASVs counts, wherever possible, were treated as compositional. Zeros were replaced by Bayesian-multiplicative replacement and counts transformed to centred log-ratios (CLR) using *zCompositions* v1.6.2 (Palarea-Albaladejo & Martín-Fernández 2015). Three analyses used another approach because the quantities they estimate have no compositional equivalent: the turnover and nestedness partition, which uses presence-absence data; phylogenetic redundancy, which is abundance-weighted; and the assembly-process null model, which is based on Bray-Curtis dissimilarity and phylogenetic turnover. These were run on the rarefied table. Multivariate distances were calculated as Aitchison distance, and ordination was visualized by principal component analysis of the same matrix.

Inference on effect sizes for different metrics was made with a Bayesian multilevel model fitted in *brms* v2.23.0 (Bürkner 2017) through *rstan* v2.32.7 (Carpenter *et al*. 2017; Stan Development Team 2020). Models included a random intercept for soil source and, where the question required it, a random stressor slope, so that each effect was estimated within each soil community and pooled across communities. Proportions were modelled with a beta likelihood and distances with a lognormal likelihood. Four chains of 4,000 iterations were run, half discarded as warmup; where divergent transitions occurred, the model was refitted with a higher target acceptance rate. Convergence was assessed with R-hat and effective sample size and is reported for every model in **Supplementary Table S13**. Effects are reported as posterior means with 95% credible intervals, and described as credible when the interval excludes zero.

Permutational multivariate analysis of variance and tests of multivariate dispersion homogeneity were computed with *vegan* v2.7-5 (Dixon 2003) using 999 permutations, fitting each term marginally as well as sequentially. Because a PERMANOVA R^2^ cannot be compared across terms of very different degrees of freedom, variance was additionally decomposed in a Bayesian multilevel model of the five leading Aitchison principal components, each standardized, with random intercepts for soil source, stressor, genotype and their pairwise interactions. Each component share of variance was computed within each posterior draw and averaged across components, weighted by the Aitchison variance each carries.

Sørensen dissimilarity was partitioned into turnover and nestedness components with betapart v1.6.1 (Baselga & Orme 2012). Because every plant appears in many pairs, pairwise values were summarized as one value per plant: the mean over other plants sharing its inoculum, stressor and genotype, and the mean over plants of every other inoculum with stressor and genotype held constant. Aitchison distances were summarized in the same way. Phylogenetic redundancy was calculated using *SYNCSA* v1.3.5 (Debastiani & Pillar 2012), from a cophenetic distance matrix derived from the ASV phylogeny with *ape* v5.8-1 (Paradis & Schliep 2019), and was modelled with observed richness as a covariate.

Differentially abundant taxa were identified with *ALDEx2* v1.36.0 (Fernandes *et al*. 2014), run separately within each soil source and contrasting each stressor against the shared control with genotypes pooled, giving one standardized effect per ASV per soil source. These were combined in a hierarchical model, effect ~ 1 + (1 | ASV) + (1 | soil source), in which the ASV term estimates the response pooled across communities. Because each ASV is estimated once per soil source, an explicit ASV-by-source interaction is not separable from the residual, and the residual is reported as that quantity. A taxon was described as a general responder when its pooled effect was credible and it moved in the same direction in at least 70% of the soil sources in which it was testable.

Turnover between every pair of plants was apportioned among heterogeneous selection, homogeneous selection, dispersal limitation, homogenizing dispersal, and drift and other processes using *iCAMP* v1.5.12 (Ning *et al*. 2020), with 1,000 randomizations and a minimum of 12 ASVs per phylogenetic bin. *iCAMP* was run separately within each soil source, so that assembly is assessed among plants that received the same starting community, and the resulting per-pair compositions were summarized per plant as above.

Co-occurrence networks were inferred with *SpiecEasi* v1.1.3 (Kurtz *et al*. 2015) on the 200 most prevalent ASVs of each group and requiring at least 14 samples. Networks were built per soil source, per soil source and stressor, and per soil source and genotype. Topology and robustness were computed with *igraph* v2.3.3 (Antonov *et al*. 2023): nodes were removed at random and in order of decreasing degree, tracking the size of the largest connected component and natural connectivity, and summarizing each curve by its area. Because robustness increases with the number of edges recovered for reasons unrelated to biology, edge count was included as a covariate in every model of robustness.

Finally, the per-plant metrics were combined in a Bayesian structural equation model, fitted as a multivariate *brms* model with a shared random intercept for soil source across equations. All variables were placed on an unbounded scale and standardized, so that path coefficients are comparable and their products interpretable as indirect effects; the coefficients of this model are therefore not comparable in magnitude with the single-equation estimates reported elsewhere. Variables that are arithmetic functions of one another were not entered together. Indirect effects were computed by enumerating every directed route through the model and multiplying coefficients within each posterior draw, so that the resulting distributions carry the uncertainty of every path they contain.

## Author contributions

Conceptualization: EF, AM; Methodology: EF, MMA, AM; Investigation: EF, MMA, AM; Visualization: AM; Writing-original draft: EF, AM; Writing-review and editing: all co-authors.

## Data availability

Data is available on NCBI SRA under BioProject PRJNA1521537, and the code is available at https://github.com/malacrino-lab/PRIN_tomato.

