## Supplementary materials for "Foliar pathogen and drought impose reproducible but community-dependent influence on the root microbiome"

### Supplementary figures

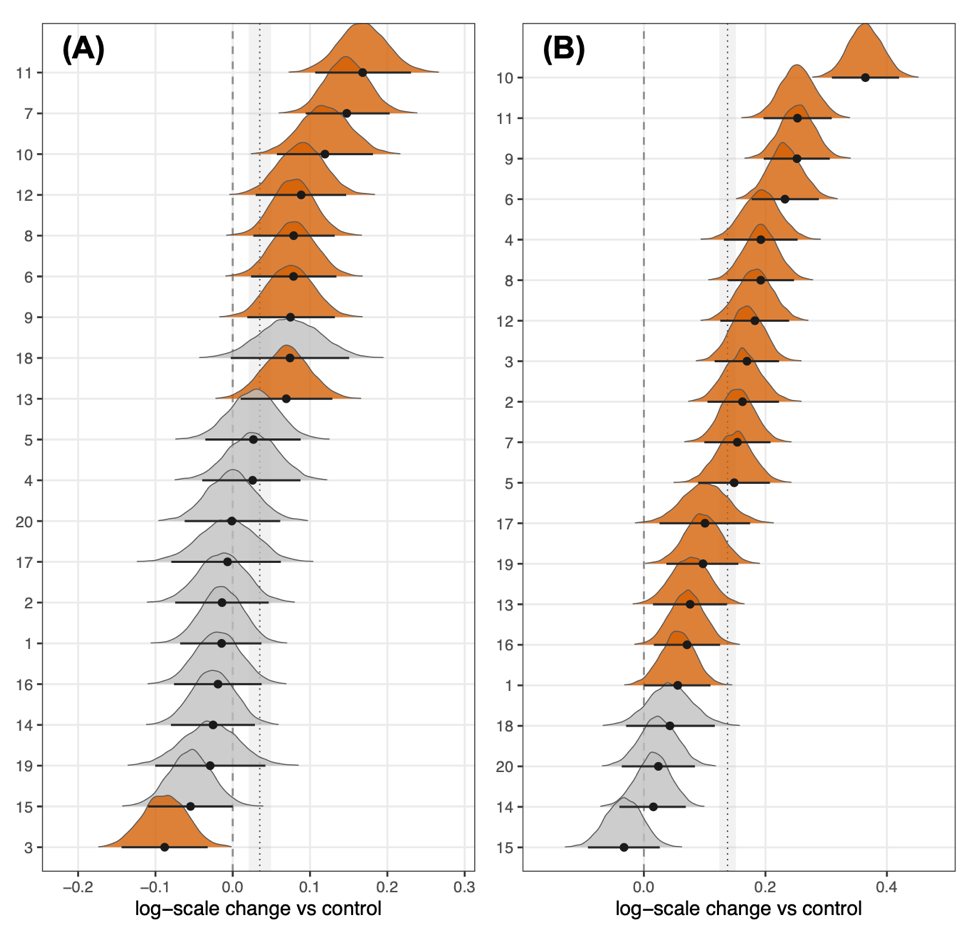

**Figure S1**. Posterior distributions of effects on microbiota composition variability under drought **(A)** and **(B)** pathogen across inocula. Densities are colored where the 95% credible interval excludes zero, the grey band marks the mean across sources, and points with bars give posterior means and 95% credible intervals.

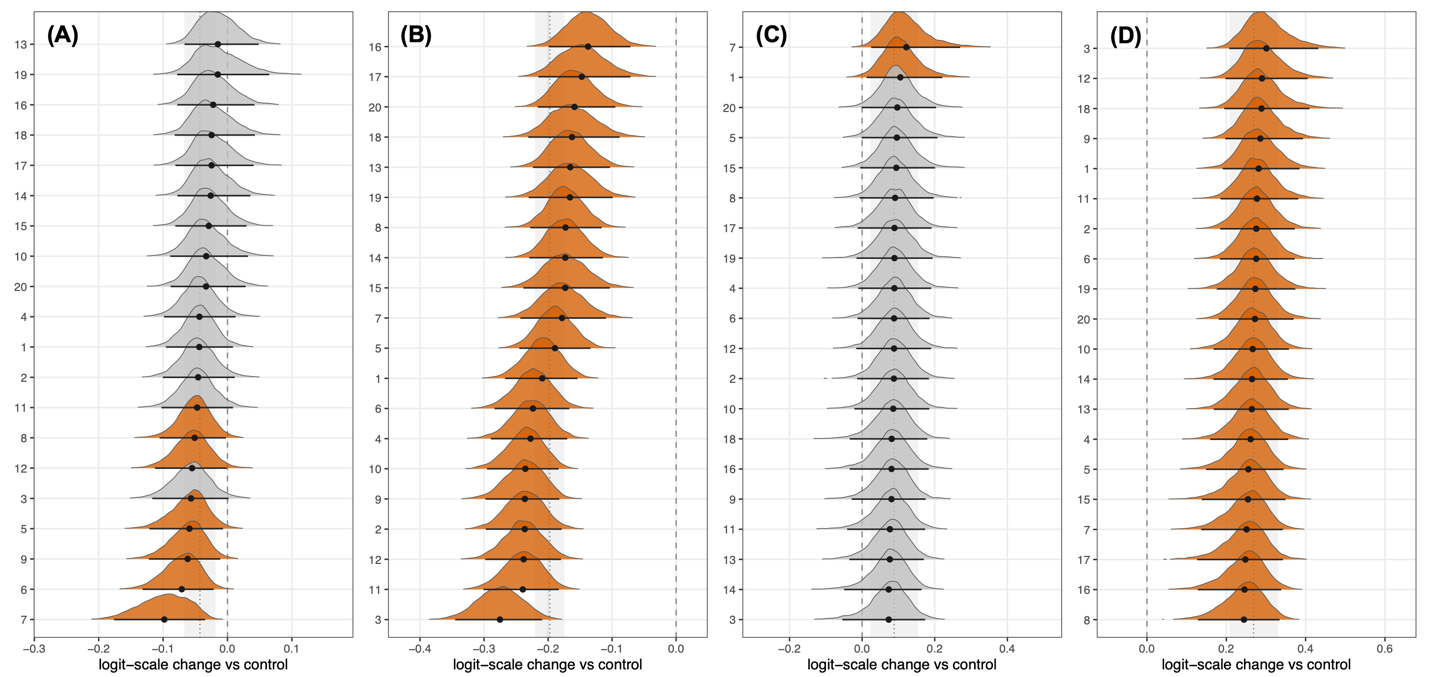

**Figure S2**. Posterior distributions of effects on microbiota turnover **(A-B)** and nestedness **(C-D)** under drought **(A,C)** and **(B,D)** pathogen across inocula. Densities are colored where the 95% credible interval excludes zero, the grey band marks the mean across sources, and points with bars give posterior means and 95% credible intervals.

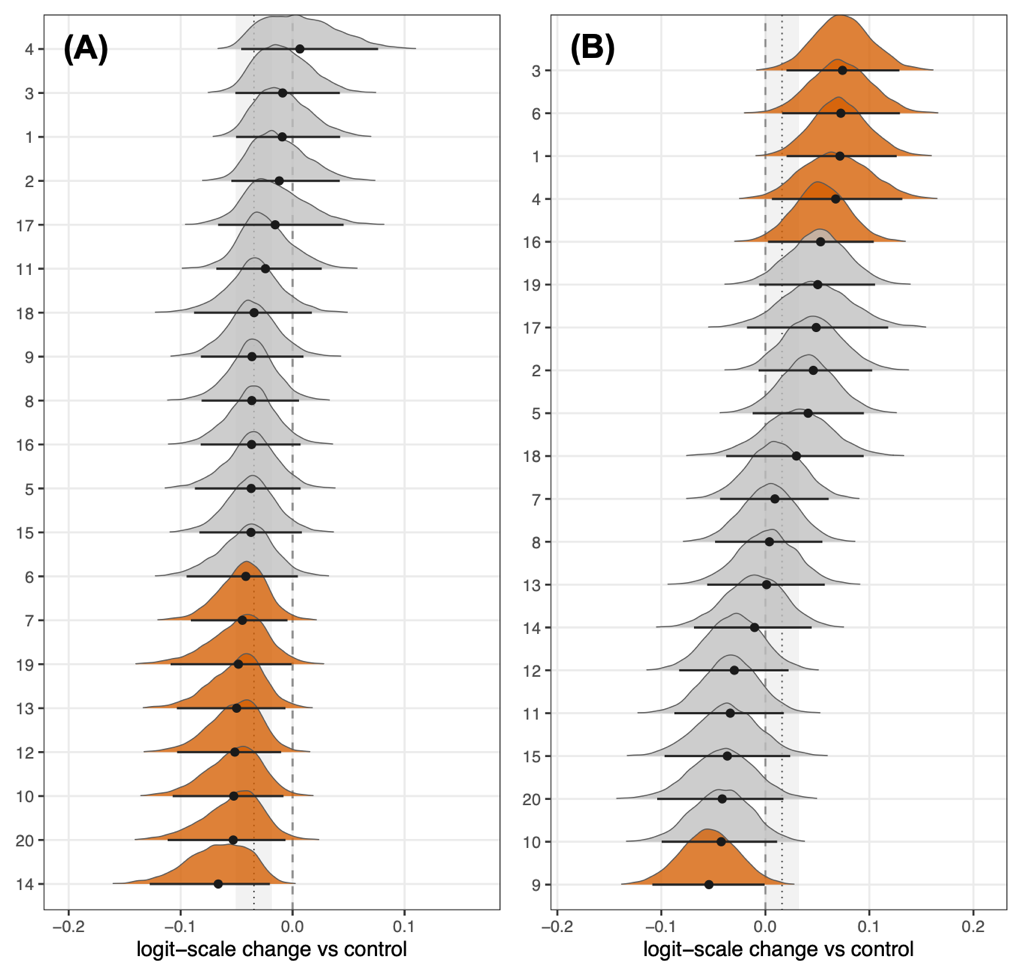

**Figure S3**. Posterior distributions of effects on phylogenetic redundancy under drought **(A)** and **(B)** pathogen across inocula. Densities are colored where the 95% credible interval excludes zero, the grey band marks the mean across sources, and points with bars give posterior means and 95% credible intervals.

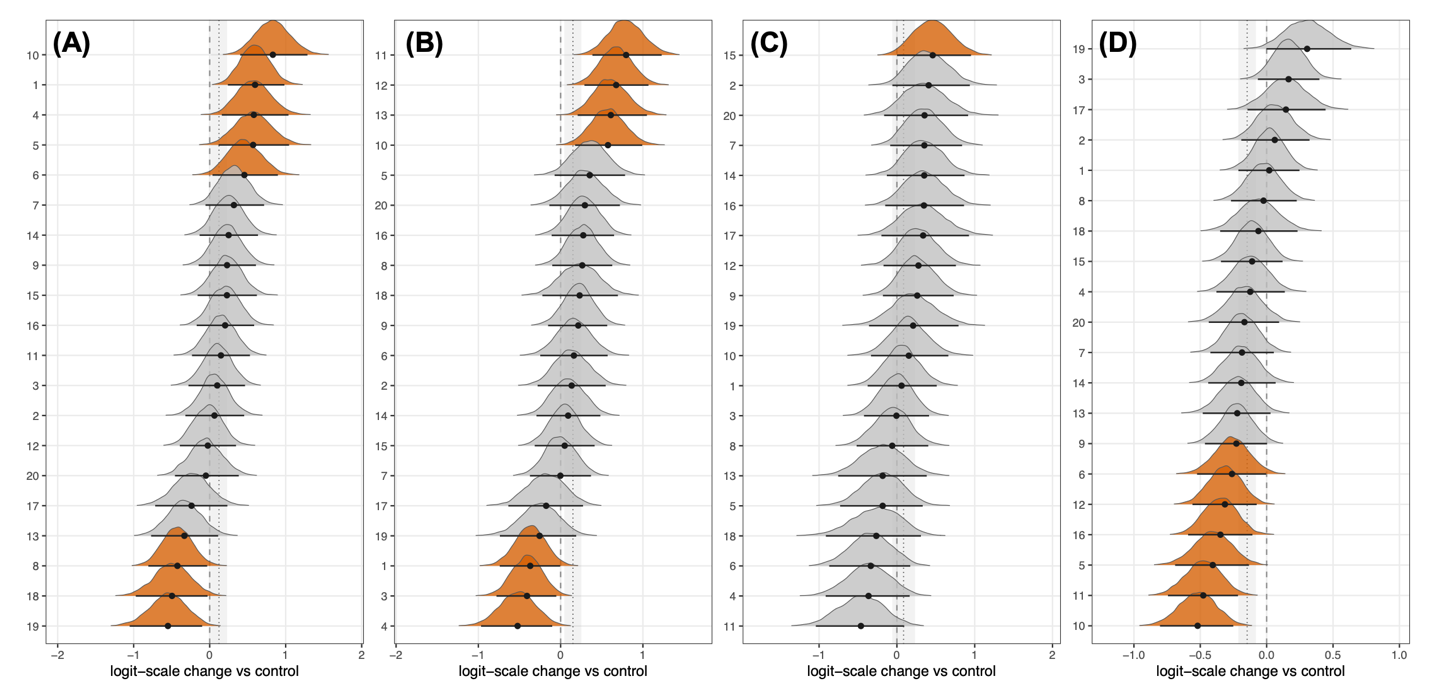

**Figure S4**. Per-inoculum effects of drought on each of the community assembly processes: **(A)** heterogeneous and homogeneous selection, **(B)** dispersal limitation, **(C)** homogenizing dispersal, **(D)** drift and others. Densities are colored where the 95% credible interval excludes zero, the grey band marks the mean across sources, and points with bars give posterior means and 95% credible intervals.

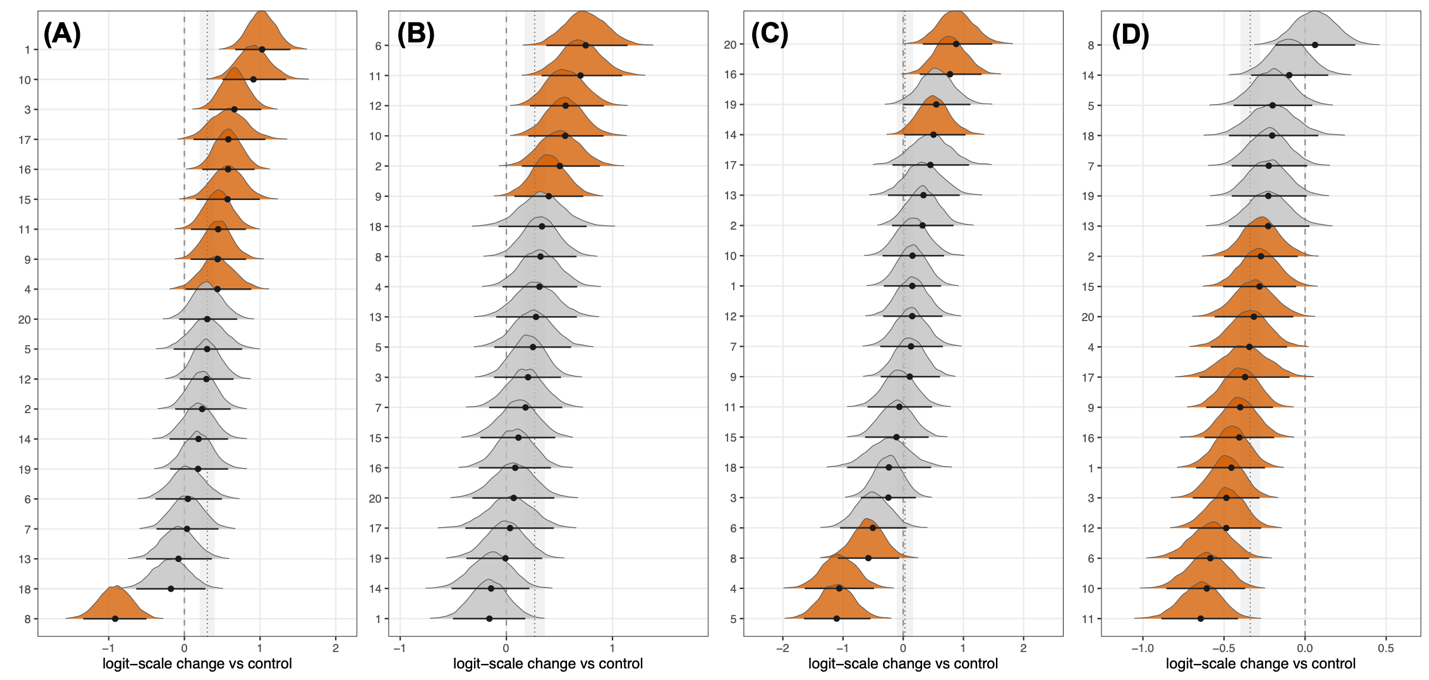

**Figure S5**. Per-inoculum effects of pathogen on each of the community assembly processes: **(A)** heterogeneous and homogeneous selection, **(B)** dispersal limitation, **(C)** homogenizing dispersal, **(D)** drift and others. Densities are colored where the 95% credible interval excludes zero, the grey band marks the mean across sources, and points with bars give posterior means and 95% credible intervals.

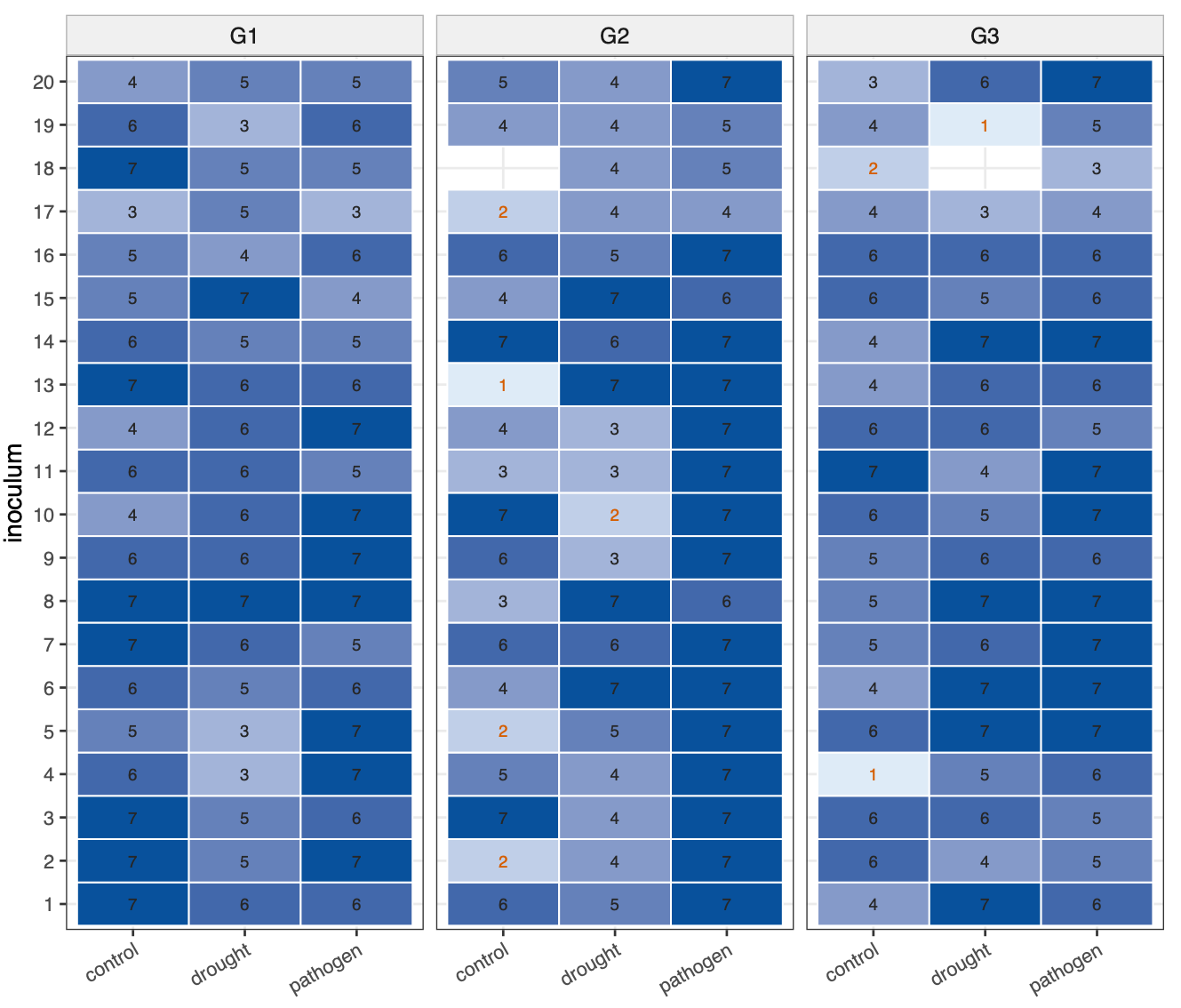

**Figure S6**. Occupancy of the experimental design after quality control. Each tile is one combination of soil inoculum (rows), stressor treatment (columns) and tomato genotype (panels), and gives the number of plants retained in that combination after quality filtering; tile shading is proportional to that number. Counts printed in orange mark combinations retaining fewer than three plants, the minimum required for a combination to contribute to models estimating combination-specific quantities. Combinations that retained no plants are not drawn. Of the 180 combinations in the full 20 × 3 × 3 design, 178 were occupied and 170 retained at least three plants, with a median of six.

### Supplementary tables

**Table S1**. Permutational multivariate analysis of variance on Aitchison distances. Marginal fits of each design factor and the full sequential model including two-way interactions; 999 permutations.

| **term** | **df** | **R^2^** | **F** | **p** | **fit** |
| --- | --- | --- | --- | --- | --- |
| inoculum | 19 | 0.0411 | 2.12 | 0.001 | marginal |
| treatment | 2 | 0.0271 | 13.35 | 0.001 | marginal |
| genotype | 2 | 0.0098 | 4.76 | 0.001 | marginal |
| inoculum | 19 | 0.0411 | 2.34 | 0.001 | sequential |
| treatment | 2 | 0.0273 | 14.79 | 0.001 | sequential |
| genotype | 2 | 0.0096 | 5.19 | 0.001 | sequential |
| inoculum:treatment | 38 | 0.073 | 2.08 | 0.001 | sequential |
| inoculum:genotype | 38 | 0.0483 | 1.38 | 0.001 | sequential |
| treatment:genotype | 4 | 0.0087 | 2.36 | 0.001 | sequential |
| Residual | 858 | 0.792 |  |  | sequential |
| Total | 961 | 1 |  |  | sequential |

**Table S2**. Variance in Aitchison space attributable to each design factor. Posterior mean share and 95% credible interval from a Bayesian multilevel model of PC1-PC5, weighted by the variance each component carries, with pairwise rank probabilities.

| **component** | **share** | **Q2.5** | **Q97.5** |
| --- | --- | --- | --- |
| residual | 0.3901 | 0.257 | 0.5096 |
| treatment | 0.297 | 0.1285 | 0.506 |
| inoculum.treatment | 0.1303 | 0.0862 | 0.1775 |
| genotype | 0.0884 | 0.016 | 0.2407 |
| treatment.genotype | 0.0415 | 0.0122 | 0.115 |
| inoculum.genotype | 0.0333 | 0.0188 | 0.0516 |
| inoculum | 0.0195 | 0.0037 | 0.0461 |

**Table S3**. Stressor and genotype effects on microbiome variability. Posterior estimates from lognormal multilevel models of variability among replicate plants and of distance between soil sources, with random stressor slopes per soil source.

| **response** | **term** | **Estimate** | **Est.Error** | **Q2.5** | **Q97.5** |
| --- | --- | --- | --- | --- | --- |
| variability among replicate plants | Intercept | 3.5605 | 0.014 | 3.5328 | 3.5889 |
| variability among replicate plants | treatmentdrought | 0.0343 | 0.019 | -0.0041 | 0.0711 |
| variability among replicate plants | treatmentpathogen | 0.1365 | 0.0253 | 0.0857 | 0.1862 |
| variability among replicate plants | genotypeG2 | -0.0041 | 0.007 | -0.0178 | 0.0097 |
| variability among replicate plants | genotypeG3 | -0.033 | 0.0069 | -0.0468 | -0.0196 |
| distance between soil sources | Intercept | 3.6264 | 0.0066 | 3.6131 | 3.6397 |
| distance between soil sources | treatmentdrought | 0.0291 | 0.0078 | 0.0137 | 0.0441 |
| distance between soil sources | treatmentpathogen | 0.1483 | 0.0088 | 0.1308 | 0.1654 |
| distance between soil sources | genotypeG2 | -0.0055 | 0.0062 | -0.0175 | 0.0064 |
| distance between soil sources | genotypeG3 | -0.032 | 0.0061 | -0.0437 | -0.02 |

**Table S4**. Stressor and genotype effects on turnover and nestedness. Population-level posterior estimates from beta regressions of the two Sorensen components, computed among replicate plants and between soil sources.

| **response** | **term** | **Estimate** | **Est.Error** | **Q2.5** | **Q97.5** |
| --- | --- | --- | --- | --- | --- |
| within_turnover | Intercept | -0.488 | 0.0239 | -0.5351 | -0.4406 |
| within_turnover | treatmentdrought | 0.0069 | 0.0195 | -0.0307 | 0.0443 |
| within_turnover | treatmentpathogen | -0.1693 | 0.0298 | -0.2278 | -0.1094 |
| within_turnover | genotypeG2 | 0.033 | 0.0164 | 0.0006 | 0.065 |
| within_turnover | genotypeG3 | 0.0502 | 0.0163 | 0.0178 | 0.0822 |
| within_nestedness | Intercept | -2.3495 | 0.0677 | -2.4853 | -2.2166 |
| within_nestedness | treatmentdrought | 0.117 | 0.0707 | -0.0233 | 0.2576 |
| within_nestedness | treatmentpathogen | 0.1879 | 0.067 | 0.0568 | 0.3195 |
| within_nestedness | genotypeG2 | 0.0015 | 0.0416 | -0.0801 | 0.0835 |
| within_nestedness | genotypeG3 | -0.0347 | 0.0414 | -0.1148 | 0.0466 |
| between_turnover | Intercept | -0.2662 | 0.0119 | -0.289 | -0.2424 |
| between_turnover | treatmentdrought | -0.0429 | 0.0151 | -0.0722 | -0.0123 |
| between_turnover | treatmentpathogen | -0.1974 | 0.0168 | -0.2306 | -0.1642 |
| between_turnover | genotypeG2 | 0.0829 | 0.0117 | 0.0607 | 0.1061 |
| between_turnover | genotypeG3 | 0.0959 | 0.0114 | 0.0741 | 0.1184 |
| between_nestedness | Intercept | -2.331 | 0.0301 | -2.3908 | -2.2731 |
| between_nestedness | treatmentdrought | 0.0883 | 0.0347 | 0.0193 | 0.1565 |
| between_nestedness | treatmentpathogen | 0.2689 | 0.033 | 0.2054 | 0.3337 |
| between_nestedness | genotypeG2 | -0.1092 | 0.03 | -0.1679 | -0.0501 |
| between_nestedness | genotypeG3 | -0.086 | 0.0297 | -0.144 | -0.0275 |

**Table S5**. Stressor and genotype effects on phylogenetic redundancy, with per-source stressor slopes.

| **response** | **term** | **Estimate** | **Est.Error** | **Q2.5** | **Q97.5** |
| --- | --- | --- | --- | --- | --- |
| redundancy | Intercept | 0.709 | 0.0099 | 0.6898 | 0.7288 |
| redundancy | treatmentdrought | -0.0345 | 0.0108 | -0.0558 | -0.0132 |
| redundancy | treatmentpathogen | 0.0162 | 0.015 | -0.0134 | 0.0455 |
| redundancy | genotypeG2 | -0.0217 | 0.0078 | -0.0368 | -0.0061 |
| redundancy | genotypeG3 | -0.0323 | 0.0077 | -0.047 | -0.0168 |
| redundancy | richness_z | -0.002 | 0.0036 | -0.0091 | 0.0052 |
| Rao entropy | Intercept | -0.8698 | 0.0144 | -0.8983 | -0.8416 |
| Rao entropy | treatmentdrought | 0.0307 | 0.0127 | 0.0055 | 0.0551 |
| Rao entropy | treatmentpathogen | -0.0305 | 0.0183 | -0.0662 | 0.0058 |
| Rao entropy | genotypeG2 | 0.0245 | 0.0099 | 0.0049 | 0.0435 |
| Rao entropy | genotypeG3 | 0.0359 | 0.0098 | 0.0164 | 0.0543 |
| Rao entropy | richness_z | 0.0515 | 0.0045 | 0.0429 | 0.0603 |

**Table S6**. Community assembly processes. Posterior stressor and genotype effects on each process and on the deterministic fraction.

| **process** | **term** | **Estimate** | **Est.Error** | **Q2.5** | **Q97.5** |
| --- | --- | --- | --- | --- | --- |
| deterministic | Intercept | -2.005 | 0.1039 | -2.2132 | -1.8036 |
| deterministic | treatmentdrought | 0.1182 | 0.1113 | -0.1024 | 0.339 |
| deterministic | treatmentpathogen | 0.2999 | 0.1173 | 0.0711 | 0.5356 |
| deterministic | genotypeG2 | -0.0398 | 0.0486 | -0.1341 | 0.0545 |
| deterministic | genotypeG3 | -0.2583 | 0.05 | -0.3559 | -0.1596 |
| HeS | Intercept | -5.0557 | 0.1016 | -5.2545 | -4.8584 |
| HeS | treatmentdrought | 0.137 | 0.1383 | -0.1355 | 0.4069 |
| HeS | treatmentpathogen | 0.1327 | 0.0947 | -0.0576 | 0.3216 |
| HeS | genotypeG2 | -0.2795 | 0.0754 | -0.4271 | -0.1325 |
| HeS | genotypeG3 | -0.1358 | 0.075 | -0.2816 | 0.0123 |
| HoS | Intercept | -2.0771 | 0.115 | -2.2988 | -1.8456 |
| HoS | treatmentdrought | 0.1119 | 0.1286 | -0.1378 | 0.3611 |
| HoS | treatmentpathogen | 0.3064 | 0.126 | 0.0541 | 0.5488 |
| HoS | genotypeG2 | -0.0108 | 0.0513 | -0.1128 | 0.0879 |
| HoS | genotypeG3 | -0.2552 | 0.0522 | -0.3568 | -0.1536 |
| DL | Intercept | -1.6801 | 0.0768 | -1.8307 | -1.5298 |
| DL | treatmentdrought | 0.1484 | 0.1105 | -0.0697 | 0.3686 |
| DL | treatmentpathogen | 0.2657 | 0.0877 | 0.0894 | 0.4417 |
| DL | genotypeG2 | -0.1009 | 0.0495 | -0.1984 | -0.0022 |
| DL | genotypeG3 | 0.0078 | 0.0481 | -0.0858 | 0.1027 |
| HD | Intercept | -3.6877 | 0.1195 | -3.9282 | -3.4583 |
| HD | treatmentdrought | 0.0868 | 0.1165 | -0.139 | 0.3143 |
| HD | treatmentpathogen | 0.0285 | 0.1456 | -0.2592 | 0.3199 |
| HD | genotypeG2 | -0.0932 | 0.0692 | -0.2278 | 0.0431 |
| HD | genotypeG3 | -0.138 | 0.0688 | -0.274 | -0.0016 |
| DR | Intercept | 0.7879 | 0.0532 | 0.6847 | 0.8953 |
| DR | treatmentdrought | -0.148 | 0.0697 | -0.2871 | -0.0099 |
| DR | treatmentpathogen | -0.3385 | 0.0601 | -0.4562 | -0.2166 |
| DR | genotypeG2 | 0.1268 | 0.0309 | 0.0674 | 0.1877 |
| DR | genotypeG3 | 0.17 | 0.0311 | 0.1088 | 0.2304 |

**Table S7**. Summary statistics across the 20 networks inferred separately for each soil source.

| **metric** | **value** |
| --- | --- |
| networks | 20 |
| mean_auc_targeted | 0.1289 |
| sd_auc_targeted | 0.05 |
| min_auc_targeted | 0.0638 |
| max_auc_targeted | 0.2573 |
| mean_auc_random | 0.3059 |
| sd_auc_random | 0.0526 |
| correlation_auc_with_edges | 0.952 |
| correlation_auc_with_modularity | -0.873 |

**Table S8**. Stressor effects on network topology and robustness. Population-level posterior estimates from Bayesian beta regressions of three network properties, fitted across the 45 networks inferred for each combination of soil source and stressor, with a random intercept for soil source. Coefficients are on the logit scale relative to the unstressed control; edges_z is the number of recovered edges, standardized, included as a covariate because robustness rises with network density for reasons unrelated to biology. Est.Error is the posterior standard deviation, Q2.5 and Q97.5 bound the 95% credible interval, and credible marks intervals excluding zero (1) or including it (0).

| **response** | **term** | **Estimate** | **Est.Error** | **Q2.5** | **Q97.5** |
| --- | --- | --- | --- | --- | --- |
| robustness, targeted | Intercept | -1.477 | 0.0434 | -1.5636 | -1.391 |
| robustness, targeted | treatmentdrought | -0.09 | 0.0597 | -0.2069 | 0.0289 |
| robustness, targeted | treatmentpathogen | -0.1058 | 0.0572 | -0.2149 | 0.0083 |
| robustness, targeted | edges_z | 0.4427 | 0.0264 | 0.3916 | 0.495 |
| robustness, random | Intercept | -0.6778 | 0.0376 | -0.7506 | -0.6038 |
| robustness, random | treatmentdrought | -0.0671 | 0.054 | -0.1738 | 0.0384 |
| robustness, random | treatmentpathogen | -0.082 | 0.0491 | -0.1778 | 0.0148 |
| robustness, random | edges_z | 0.2948 | 0.022 | 0.2518 | 0.3387 |
| modularity | Intercept | 0.741 | 0.036 | 0.6705 | 0.8136 |
| modularity | treatmentdrought | 0.022 | 0.0509 | -0.0782 | 0.1235 |
| modularity | treatmentpathogen | 0.0585 | 0.0464 | -0.034 | 0.1498 |
| modularity | edges_z | -0.3014 | 0.0211 | -0.3435 | -0.2584 |

**Table S9**. Taxa responding consistently to each stressor across soil communities. Pooled posterior effect, 95% credible interval, number of soil sources in which the ASV was testable, directional consistency and taxonomy, for ASVs meeting both criteria. SEE EXCEL FILE.

**Table S10**. Cross-inoculum summary of stressor effects for every ASV, giving the mean and standard deviation of the variance components of the hierarchical meta-analysis.

| **stressor** | **component** | **sd** | **Q2.5** | **Q97.5** |
| --- | --- | --- | --- | --- |
| drought | ASV | 0.2519 | 0.2231 | 0.2849 |
| drought | inoculum | 0.0108 | 0.0005 | 0.0278 |
| drought | ASV:inoculum (residual) | 0.2981 | 0.2904 | 0.3064 |
| pathogen | ASV | 0.2467 | 0.2183 | 0.2785 |
| pathogen | inoculum | 0.025 | 0.004 | 0.0474 |
| pathogen | ASV:inoculum (residual) | 0.337 | 0.3282 | 0.3463 |

**Table S11**. Variance explained by each equation within the structural equation model. Bayesian R2 per equation

| **equation** | **Estimate** | **Est.Error** | **Q2.5** | **Q97.5** |
| --- | --- | --- | --- | --- |
| R2det | 0.1192 | 0.0201 | 0.08 | 0.1603 |
| R2tur | 0.3482 | 0.0212 | 0.3058 | 0.3882 |
| R2nes | 0.1152 | 0.0196 | 0.0774 | 0.1545 |
| R2red | 0.123 | 0.0205 | 0.0844 | 0.1645 |
| R2vrb | 0.4835 | 0.0179 | 0.4476 | 0.5174 |

**Table S12**. Standardized path coefficients of the structural equation model. All modelled paths with posterior means and 95% credible intervals, and Bayesian R2 per equation. SEE EXCEL FILE.

**Table S13**. Source of soil inocula. For field margin and uncultivated field there was not a dominant plant species to report.

| **Inoculum** | **Type** | **Plant species** |
| --- | --- | --- |
| 1 | agricultural | fava bean |
| 2 | agricultural | no crop present |
| 3 | agricultural | cherry |
| 4 | field margin |  |
| 5 | agricultural | apple |
| 6 | agricultural | cherry |
| 7 | field margin |  |
| 8 | agricultural | hazelnut |
| 9 | agricultural | strawberry |
| 10 | field margin |  |
| 11 | field margin |  |
| 12 | field margin |  |
| 13 | agricultural | chestnut |
| 14 | agricultural | oat |
| 15 | agricultural | wheat |
| 16 | field margin |  |
| 17 | uncultivated field |  |
| 18 | forest | pine |
| 19 | forest | oak |
| 20 | forest | oak |

**Table S14**. Convergence diagnostics for all fitted Bayesian models.

| **model** | **max_rhat** | **min_bulk_ess** | **min_tail_ess** | **divergences** |
| --- | --- | --- | --- | --- |
| between-inocula distance | 1.0083 | 916 | 664 | 0 |
| beta_within_turnover | 1.0042 | 876 | 1891 | 0 |
| iCAMP HeS | 1.0041 | 1089 | 1651 | 0 |
| phylogenetic redundancy | 1.004 | 1130 | 1178 | 0 |
| beta_between_turnover | 1.0034 | 1050 | 1268 | 0 |
| iCAMP HoS | 1.0033 | 1735 | 2621 | 0 |
| Rao entropy | 1.0032 | 1158 | 1666 | 0 |
| within-cell variability | 1.0031 | 1701 | 3268 | 0 |
| ALDEx2 meta-analysis drought | 1.0031 | 1161 | 2193 | 0 |
| iCAMP HD | 1.0028 | 1481 | 2041 | 0 |
| ALDEx2 meta-analysis pathogen | 1.0025 | 1445 | 2218 | 0 |
| PC contrasts PC1 | 1.0022 | 1935 | 3231 | 0 |
| beta_within_nestedness | 1.0021 | 1406 | 1452 | 0 |
| iCAMP deterministic fraction | 1.0021 | 1694 | 2728 | 0 |
| iCAMP DR | 1.0021 | 1705 | 3464 | 0 |
| variance decomposition PC5 | 1.002 | 2384 | 4899 | 0 |
| PC contrasts PC2 | 1.0019 | 1543 | 2877 | 0 |
| beta_between_nestedness | 1.0018 | 2066 | 3130 | 0 |
| network robustness, targeted attack | 1.0017 | 2654 | 3908 | 0 |
| network robustness, random attack | 1.0016 | 2680 | 3511 | 0 |
| network robustness by genotype | 1.0016 | 2676 | 3716 | 0 |
| PC contrasts PC3 | 1.0015 | 1877 | 3675 | 0 |
| variance decomposition PC2 | 1.0013 | 2662 | 5777 | 0 |
| variance decomposition PC4 | 1.0013 | 2062 | 3512 | 0 |
| iCAMP DL | 1.0013 | 1779 | 3159 | 0 |
| variance decomposition PC3 | 1.0012 | 3453 | 5755 | 0 |
| variance decomposition PC1 | 1.001 | 2744 | 5386 | 0 |
| network modularity | 1.0009 | 2974 | 3785 | 0 |
